# Efficacy of fecal sampling as a gut proxy in the study of chicken gut microbiota

**DOI:** 10.1101/313577

**Authors:** Wei Yan, Jiangxia Zheng, Chaoliang Wen, Congliang Ji, Dexiang Zhang, Yonghua Chen, Congjiao Sun, Ning Yang

## Abstract

**Background:** Despite the convenience and noninvasiveness of fecal sampling, the fecal microbiota does not fully represent that of the gastrointestinal (GI) tract, and the efficacy of fecal sampling to accurately represent the gut microbiota in birds is poorly understood. In this study, we aim to identify the efficacy of feces as a gut proxy in birds using chickens as a model. We collected 1,026 samples from 206 chickens, including duodenum, jejunum, ileum, cecum and feces samples, for 16S rRNA amplicon sequencing analyses.

**Results:** In this study, the efficacy of feces as a gut proxy was partitioned to microbial community membership and community structure. Most taxa in the small intestine (84.11 – 87.28%) and ceca (99.39%) could be identified in feces. Microbial community membership was reflected with a gut anatomic feature, but community structure was not. Excluding shared microbes, the small intestine and ceca contributed 34.12 and 5.83% of the total fecal members, respectively. The composition of Firmicutes members in the small intestine and that of Actinobacteria, Bacteroidetes, Firmicutes and Proteobacteria members in the ceca could be well mirrored by the observations in fecal samples (*ρ* = 0.54 – 0.71 and 0.71 – 0.78, respectively, *P* < 0.001). However, there were few significant correlations for each genus between feces and each of the 4 gut segments, and these correlations were not high (ρ = −0.2 – 0.4, *P* < 0.05) for most genera.

**Conclusions:** Our results provide evidence that the good potential of feces to identify most taxa in chicken guts, but it should be interpreted with caution by using feces as a proxy for gut in microbial structure analyses. This work provides insights and future directions regarding the usage of fecal samples in studies of the gut microbiome.

## Background

Many studies have reported the important roles of gut microbiota in host metabolism and health in humans [1], other mammals [2] and birds [3]. Because of the convenience and noninvasiveness of fecal sampling, most studies use fecal samples as a proxy to study the gut microbiota, despite the increasing recognition that fecal microbial populations may not be fully representative of those in the contents or mucosa of the gastrointestinal (GI) tract [4, 5]. Therefore, a comprehensive understanding of the efficacy of using fecal samples as a proxy to study the GI microbiota would help improve longitudinal analyses of microbiota and the application of fecal samples [6, 7].

Among birds, the chicken is frequently used as a research model, and its GI microbiota has been studied previously [8-12]. In several studies, the microbiota present in different GI segments have been investigated using traditional sequencing methods [13] or high-throughput sequencing techniques [14, 15]. However, these studies had small sample sizes (N = 3 – 8), were primarily aimed at examining the spatial heterogeneity among different segments and did not focus on the spatial microbiota relationships between feces and the GI tract.

Compared with most mammals, the cecum in birds has been reported to play important roles in metabolism, such as in the digestion of cellulose, starch and other resistant polysaccharides [16, 17] and in the absorption of nutrients [18] and water [19]. Microbial compositions and functions in chicken ceca have been reported in many studies [20, 21]. In addition, Stanley et al. [22] examined the microbial relationships between the ceca and feces and observed that 88.55% of all operational taxonomic units (OTUs) were shared. However, the microbial relationships between the ceca and small intestine (including the duodenum, jejunum and ileum), which would help provide an integrated view of gut microbial relationships, were rarely reported.

Here, we performed large-scale sequencing surveys and focused on the efficacy of using feces to represent the GI microbiota in chickens. The efficacy was partitioned into microbial community membership and structure to gain a comprehensive view to improve our understanding of the efficacy of the use of feces as a proxy to study the gut microbiota and their spatial relationships in the gut.

## Results

### Sequencing data

The 16S rRNA gene-based sequencing from 206 chickens produced 62,193,309 reads, 58,959,487 of which remained after quality filtration. The average number of sequences per sample was 57,465 and the number of sequences per sample ranged from 22,321 to 224,188.

### Landscape and quantification of microbial relationships among feces, ceca and small intestine

To gain an overview of the microbial relationships among the chicken duodenum, jejunum, ileum, ceca and feces, unweighted UniFrac distances (community membership; presence/absence of taxa) and weighted UniFrac distances (community structure; taking the relative abundances of taxa into account) were used to perform principal coordinates analysis (PCoA; Fig. 1A, B). The variation in community memberships among different sites were primarily explained by the sites origin (Fig. 1A), but the community structures showed both the sites origin and interindividual variation (Fig. 1B). In particular, the cecal microbial community exhibited a distant relationship with the small intestine community, and the microbial community in feces showed an intermediate relationship between those of the ceca and small intestines.

**Fig. 1.**
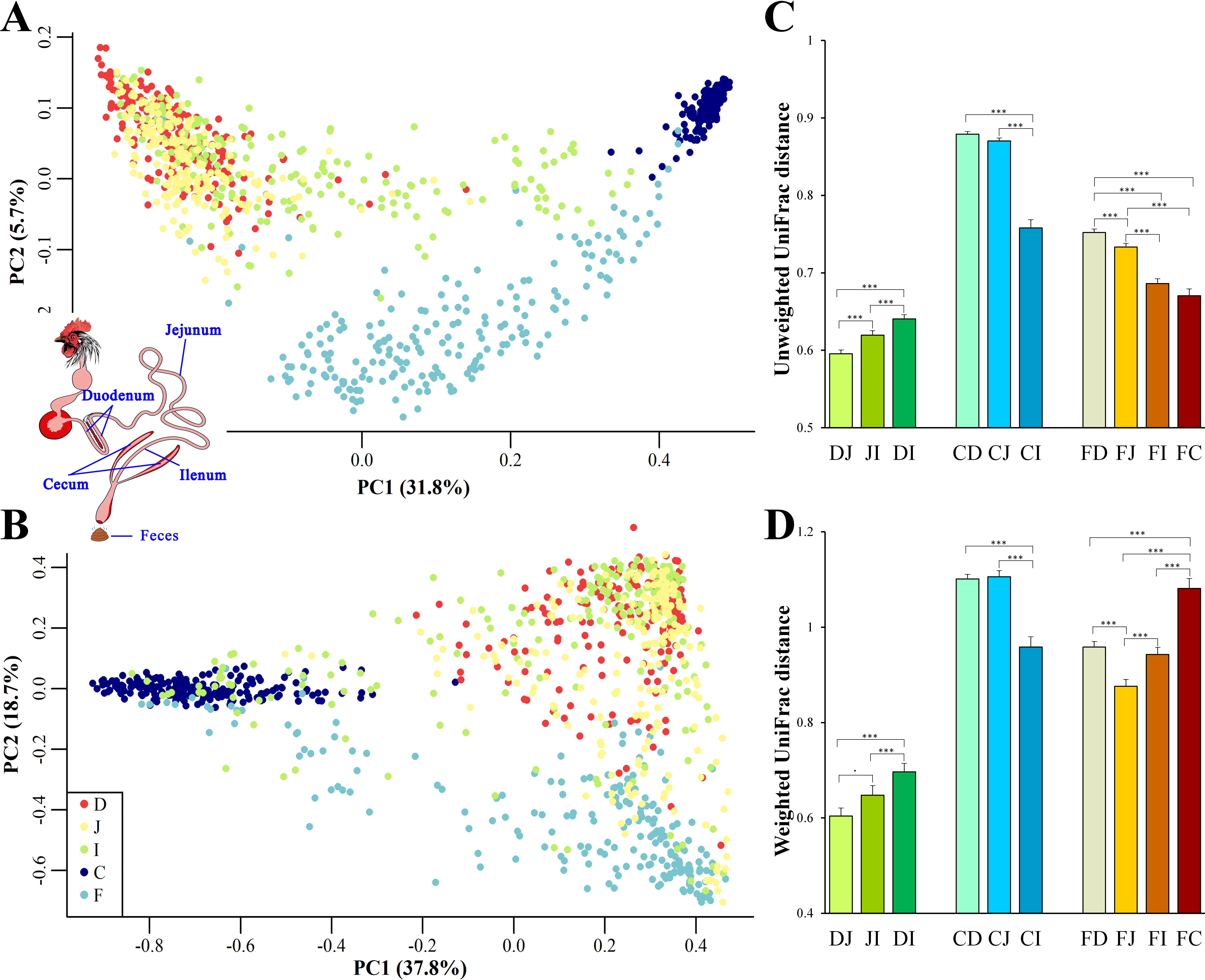
Site origin and inter-individual effects on the shape of microbial community membership and structure. (A) Principal coordinates analysis (PCoA) with unweighted UniFrac distance. Each dot represents a sample from duodenum (D), jejunum (J), ileum (I), cecum (C) or feces (F). PC1 and PC2 represent the top two principal coordinates that captured the most variation, with the fraction of variation captured by that coordinate shown as a percent. (B) PCoA plot with weighted UniFrac distance, similar to (A). (C) Unweighted UniFrac distance (mean ± SEM) between two sampling sites. DJ represents the UniFrac distance between the duodenal and jejunal microbial community, and it was the same as DI, JI, CD, CJ, CI, FD, FJ, FI and FC. Asterisks indicate the significance of the paired *t*-test: \*\*\**P* < 0.001, \*\**P* < 0.01, \**P* < 0.05 and .*P* < 0.1. (D) Weighted UniFrac distance between two sampling sites, similar to (C).

UniFrac distances between two samples from all assayed sites within each individual were calculated to quantify the spatial relationships of the gut microbiota. When the community membership was considered alone, the UniFrac distance decreased along the gut anatomical locations from the farthest to the nearest sites between fecal and duodenal, jejunal, ileal or cecal samples (FD, FJ, FI or FC, respectively, in Fig. 1C), presenting clear anatomical differences. However, when taking the community structure into account, the UniFrac distance increased in FI and FC compared with that in FJ (Fig. 1D). This finding might be explained by the exchange of contents between the ileum and ceca, suggesting that the specific cecal microbial structure influences the microbial communities in the ileum and feces.

Among all pairs, the unweighted UniFrac distance between the cecal and duodenal as well as jejunum samples were highest (*P* < 0.05), and that between duodenal and jejunal samples was lowest (*P* < 0.05; Fig. 1C and Additional file 1-2: Table S1-S2). Regarding the weighted UniFrac distances, cecal samples had similar distances to the duodenal and jejunal samples, and these distances were greater than for the other pairs (*P* < 0.05), whereas the lowest distance was observed between duodenal and jejunal samples (*P* < 0.1; Fig. 1D and Additional file 1-2: Table S1-S2). These results suggest that limited differences exist within small intestinal microbial communities, while the microbial structure in the ceca is quite distinct from those in the small intestine.

### Analyses of shared and exclusive microbial members

Given that both community membership and structure influence the microbial relationships among the feces, ceca and small intestine, we next evaluated the extent to which the spatial relationships were influenced by the above two factors. The shared and exclusive OTUs were calculated to assess the influence of the microbial community membership. To decrease the data noise, only OTUs present in more than 3 samples at each sampling site were used to analyze the effect of microbial membership. We observed that 971 OTUs, accounting for 30.9% of the total OTUs, were shared across all sites (Fig. 2A), and these shared OTUs can be referred to as the “core” microbiota in the gut. These OTUs represented different proportions of sequences in different sites and were especially high in fecal samples (96.50%; Fig. 2B), indicating that the most abundant members detected in fecal samples belonged to these “core” microbiota. At the genus level, these core taxa were primarily classified as *Bacteroides, Intestinibacter, Lactobacillus, Rikenellaceae RC9 gut group* and *Gallibacterium* (Additional file 3: Figure S1A). It is noteworthy that 5.88% of the “core” microbiota sequences were not assigned and that most of these sequences (71.40%) were detected in the cecal samples (small pie chart in Additional file 3: Figure S1A), suggesting that most of these unassigned taxa tended to be anaerobic microbes.

**Fig. 2.**
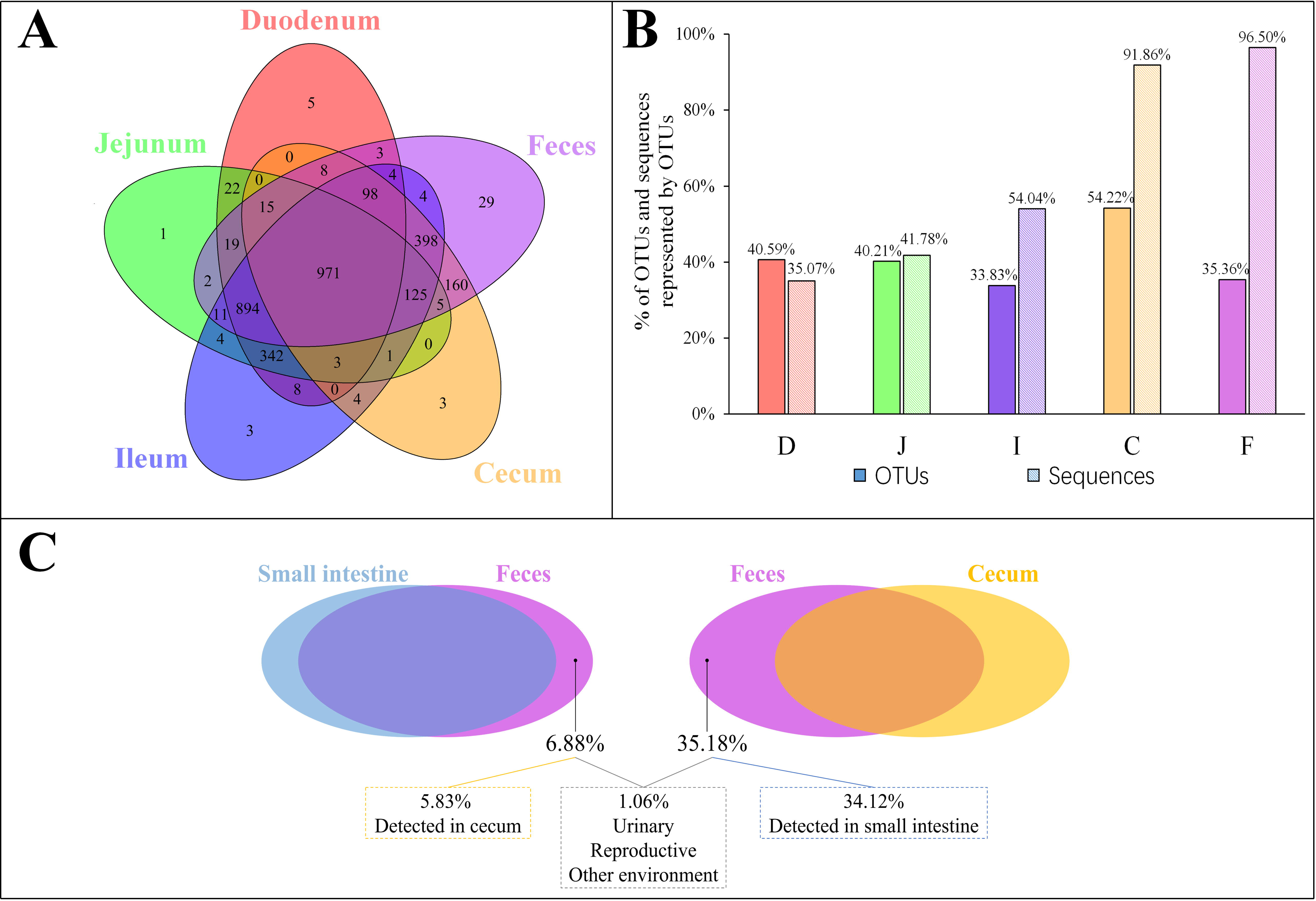
OTUs shared across different sampling sites. (A) Venn diagram demonstrating that the taxa overlap among different sampling sites. (B) The percentage of core OTUs and sequences represented by these OTUs in the duodenal (D), jejunal (J), ileal (I), cecal (C) and fecal (F) samples. (C) The percentage of OTUs in feces exclusively contributed by small intestine or cecum, and the percentage of OTUs in feces was below the limit of detection in the gastrointestinal tract.

Most OTUs in the small intestine (84.11 – 87.28%) and cecal (99.39%) samples could be identified as fecal OTUs (Table 1), indicating that feces would be a good proxy for identifying species in the gut microbiota. However, some OTUs that were present in the GI tract (12.72 – 15.89% in small intestinal and 0.61% in cecal samples) remained undetected in fecal samples (Table 1) and members of *Clostridiales, Rhizobiales, Xanthomonadales* and *Bacteroidales* appeared to be particularly undetected in feces (Additional file 4: Table S3).

**Table 1.** Shared and exclusive OTUs between two of sampling sites

| Site1 | Site2 | Shared OTUs |  | Exclusive OTUs |  |
| --- | --- | --- | --- | --- | --- |
|  |  | In site1, % | In site2, % | In site1, % | In site2, % |
| D<sup>1</sup> | J | 94.73<sup>2</sup> | 93.83 | 5.27 | 6.17 |
|  |  | 99.86<sup>3</sup> | 99.94 | 0.14 | 0.06 |
| D | I | 96.99 | 80.84 | 3.01 | 19.16 |
|  |  | 99.57 | 99.28 | 0.43 | 0.72 |
| J | I | 97.35 | 81.92 | 2.65 | 18.08 |
|  |  | 99.79 | 99.48 | 0.21 | 0.52 |
| C | D | 61.14 | 45.78 | 38.86 | 54.22 |
|  |  | 92.78 | 35.19 | 7.22 | 64.81 |
| C | J | 62.53 | 46.38 | 37.47 | 53.62 |
|  |  | 93.28 | 41.86 | 6.72 | 58.14 |
| C | I | 89.34 | 55.75 | 10.66 | 44.25 |
|  |  | 99.17 | 54.75 | 0.83 | 45.25 |
| F | D | 73.27 | 84.11 | 26.73 | 15.89 |
|  |  | 98.57 | 96.94 | 1.43 | 3.06 |
| F | J | 74.36 | 84.55 | 25.64 | 15.45 |
|  |  | 98.69 | 97.84 | 1.31 | 2.16 |
| F | I | 91.22 | 87.28 | 8.78 | 12.72 |
|  |  | 99.23 | 97.02 | 0.77 | 2.98 |
| F | C | 64.82 | 99.39 | 35.18 | 0.61 |
|  |  | 98.13 | 99.96 | 1.87 | 0.04 |

<sup>1</sup>D, J, I, C and F denotes the microbial community of duodenum, jejunum, ileum, cecum and feces, respectively. <sup>2</sup>The percentage of shared or exclusive OTUs; <sup>3</sup>The percentage of sequences shared or exclusive OTUs represent.

Microbial communities in the small intestine and ceca did not contribute equally to the fecal microbial members, as 35.18% of fecal OTUs were not identified in cecal samples, most of which (34.12%) could be identified in small intestinal niches (Table 1, Fig. 2C). These OTUs were primarily from the orders *Clostridiales, Lactobacillales, Pseudomonadales, Rickettsiales* and so on (Additional file 3: Figure S1B) and were considered exclusive contributors of the small intestinal microbiota to fecal microbial members. The ceca exclusively contributed 5.83% of OTUs to the observed fecal members, representing 0.28% of the fecal sample sequences and consisting of taxa primarily from the orders *Bacteroidales, Rhizobiales, Clostridiales, Micrococcales* and *Flavobacteriales* (Fig 2C and Additional file 3: Figure S1C).

### Correlation analyses of microbial abundances

Because community structure also affects the spatial relationships of gut microbiota, we next performed Spearman correlation analyses between the mean fecal and segmental genera abundance to evaluate the effects of community structure and assess the extent to which the microbial community in the GI tract was reflected in the fecal samples (Fig. 3). If a high correlation was observed between two sites, the differences in abundance between sites were considered highly consistent, so that the abundance at one site had the potential to be a good proxy for the abundance at another. The microbial composition of feces was correlated with those in the small intestine (Spearman: *ρ* = 0.38; *P* < 0.001) and in the combination of small intestine and ceca (*ρ* = 0.48; *P* < 0.001; Fig. 3). We then performed similar analyses to identify the correlation bias in predominant phyla (Actinobacteria, Bacteroidetes, Firmicutes and Proteobacteria; Additional file 5: Figure S2). Genera of the Firmicutes and Proteobacteria phyla in fecal samples showed moderate to high correlations with those at all four GI sites (*ρ* = 0.40 – 0.76, *P* < 0.001). In particular, fecal samples were well representative of Firmicutes members in both the small intestine and ceca (*ρ* = 0.54 – 0.71, *P* < 0.001) and of Actinobacteria, Bacteroidetes and Proteobacteria members in the ceca (*ρ* = 0.74 – 0.78, *P* < 0.001). However, Actinobacteria members in the small intestine might not be well represented in fecal samples (*ρ* = 0.13 – 0.22, *P* > 0.05).

**Fig. 3.**
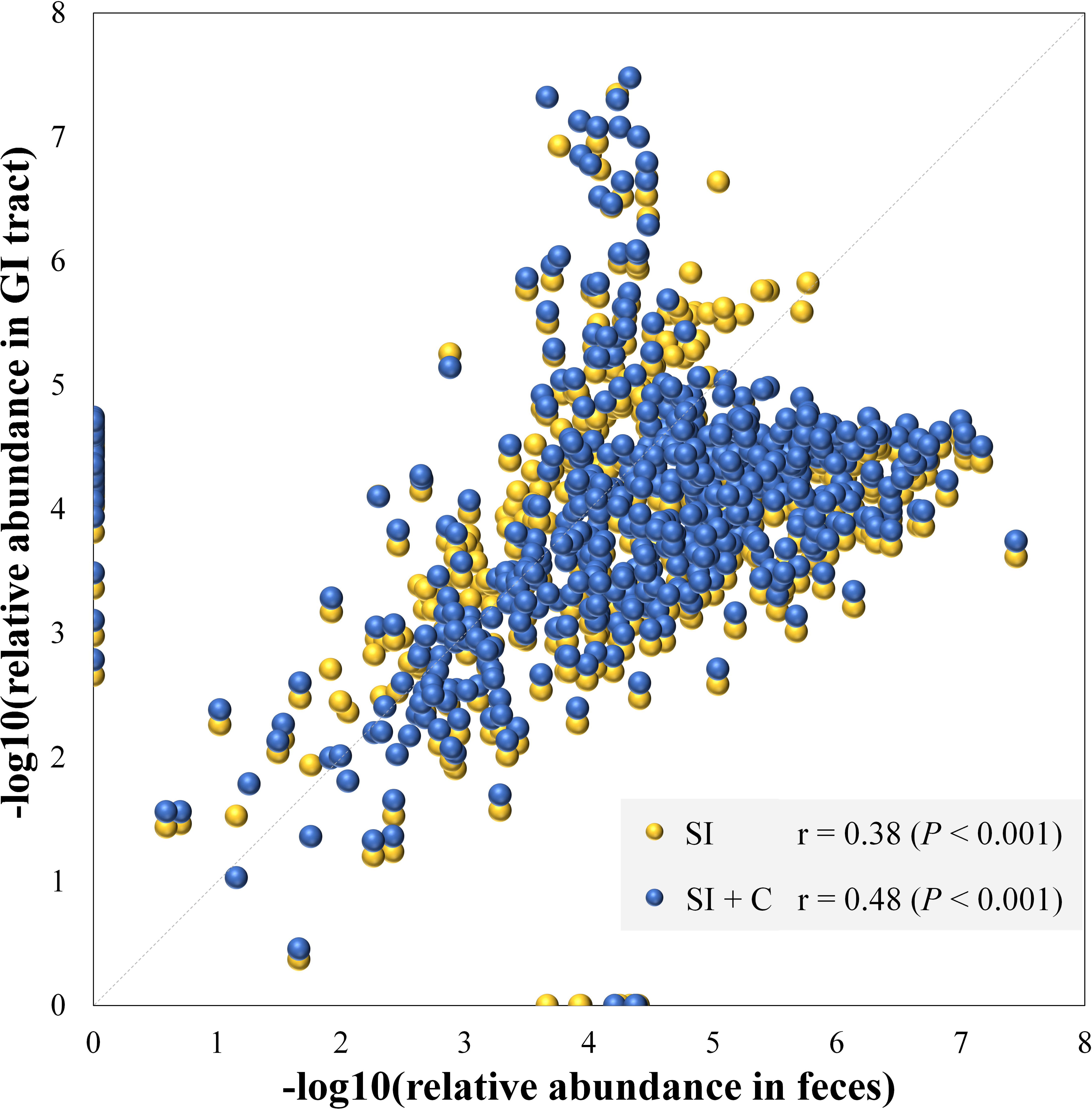
Microbial compositions in feces mirror those in the gastrointestinal tract. Each dot represents a genus. The average relative abundance of each genus in feces is transferred by negative logarithm and shown at x-axis. The average relative abundance of each genus in small intestine (SI) or intestine including small intestine and ceca (SI + C) is transferred by negative logarithm and shown at y-axis. Spearman’s rho was calculated with the negative logarithm-transferred relative abundances between feces and SI (or SI + C).

A follow-up question concerned the extent to which each microbe correlated between two sites. To address this issue, Spearman correlation tests were performed for each genus between two sites. The genera with abundances over 0.1% at either compared site with a significant correlation (*P* < 0.05) are summarized in Fig. 4 and Additional file 6: Table S4. Between the fecal and each of the 4 gut segmental samples, a limited number of significant correlations (*P* < 0.05) were observed, and these correlations were not high (*ρ* = −0.2 – 0.4, *P* < 0.05) for each genus. Most genera with significant correlations belonged to the phyla Firmicutes and Proteobacteria. However, more significant and moderate correlations were observed between two of the small intestinal segments, and most of the genera with significant correlations were also from the phyla Firmicutes and Proteobacteria (Additional file 6: Table S4). The results suggest that the gut microbiota structures could be moderately reflected by fecal samples when taking all genera into consideration simultaneously, but analyses of fluctuations in abundance for a specific genus should be interpreted with caution.

**Fig. 4.**
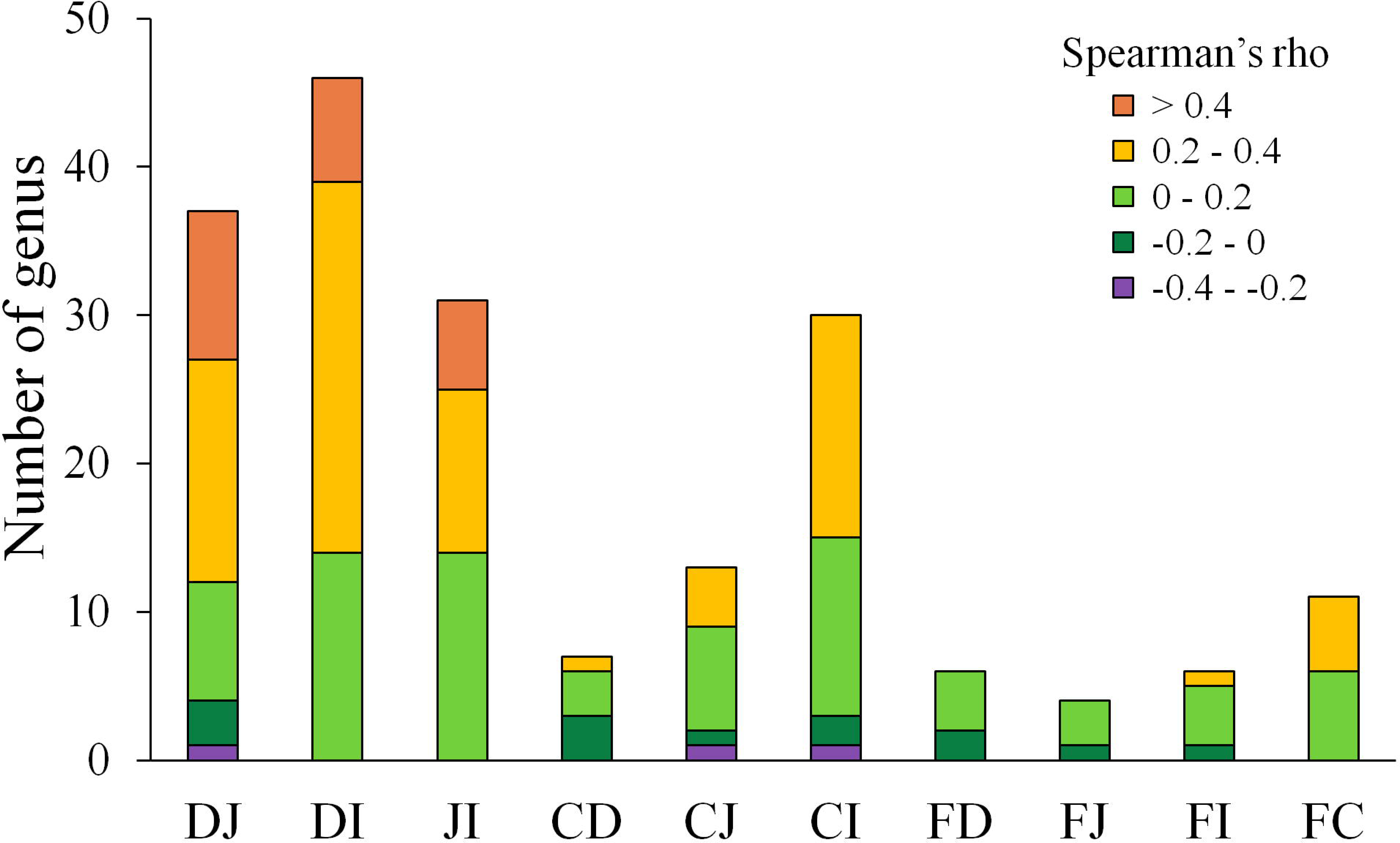
Distribution of Spearman correlations for each genus between two sites. D, J, I, C and F denote the microbial communities of the duodenum, jejunum, ileum, cecum and feces, respectively. Only genera with an abundance > 0.1% at either site of comparison and significant correlations (*P* < 0.05) are shown.

Although microbes at one site were weakly correlated with the corresponding microbes at another site, certain patterns were observed in some cases, as exemplified by the genus *Campylobacter* (Additional file 6: Table S4). The abundance of this genus in ceca exhibited consistent correlations with that observed in the jejunum (*ρ* = 0.21, *P* < 0.05) and ileum (*ρ* = 0.37, *P* < 0.05). In ileal samples, this genus was correlated with that measured in fecal samples (*ρ* = 0.19, *P* < 0.05), while no correlation was observed between cecal and fecal samples. This finding indicates that *Campylobacter* has great colonization ability in the distal gut of chickens, especially in ceca, and most *Campylobacter* contributions to the fecal composition are probably from the ileum, but not from the ceca.

## Discussion

This study is a large-scale sequencing assessment of the efficacy of using fecal samples as a proxy for the gut microbiota in birds. In this study, we comprehensively examined the community membership and structure of the chicken gut microbiome at five different biogeographic sites within 206 individual animals. We showed that fecal samples were good proxies for detecting the presence/absence of GI microbial members because most GI tract members could be detected within anatomic features in fecal samples (microbial communities in feces showed increasing similarities to those in the GI tract along the duodenum-jejunum-ileum-ceca axis). However, phyla bias and interindividual effects were observed to affect the efficacy of using fecal samples to study GI microbial abundance.

We also should note that the next-generation sequencing (NGS) approach could not absolutely detect all microbes in the gut because of some limitations of NGS method [23, 24]. Some microbes that may be present at lower levels than the limit of detection. Therefore, some OTUs that were not detected in feces but were found in the small intestine or ceca probably exist but remain below the detection limit or filtration criteria.

Similar to the current study, a high proportion of shared OTUs has been previously observed between fecal and cecal samples in chickens [22]. Similarly, a study in house mice observed that 93.3% of OTUs were shared between fecal and lower GI samples [25]. Another chicken study indicated that the GI origin is a primary determinant for the chicken fecal microbiota composition [26], supporting the high proportion of shared OTUs between feces and the four gut segments observed in the current study. These results indicate that fecal samples have good potential for identifying microbial members derived from the GI tract. However, another chicken study by Choi et al. [14] observed low percentages of shared OTUs between segments. A major reason for the differences among studies might be the small sample size in Choi’s study, which would increase the sensitivity of the results with respect to individual variation. Moreover, the presence/absence of microbial members in the GI tract was observed to be reflected by fecal samples in a given anatomical feature, i.e., fecal samples had more similarities in community membership to those in ileal and cecal samples than to those in duodenal and jejunal samples, consistent with previous reports in birds [15] and mammals [25, 27, 28].

As for microbial community structure, the efficacy of using fecal samples to represent the gut microbiota structure did not work as well as for community membership. First, the weighted UniFrac distances between feces and each of intestinal segments were significantly higher than the corresponding unweighted UniFrac distances (Additional file 7: Figure S3), suggesting that taking the abundance into account significantly increased the dissimilarity between feces and each of the GI segments. Second, the abundances of most taxa were significantly different between fecal and GI samples (Additional file 8: Table S5), consistent with previous studies[13, 15, 29]. Third, the correlations between the mean fecal and segmental genera abundances were moderate, similar to the results in rhesus macaques [30]. However, these correlations display bias among different phyla, i.e., different phyla in the GI tract are differentially mirrored by fecal samples. Fourth, significant correlations (*P* < 0.05) of each microbe between fecal and segmental samples were low and rare, suggesting that the efficacy of using fecal samples to represent microbial abundance was affected by the interindividual effect. A similar effect has also been observed in humans [4].

Previous studies in humans [4, 31] and other mammals [30, 32] have also addressed the issue of whether fecal samples are good representatives for GI microbial analyses. Although the conclusions may not be fully consistent, nearly all studies reached a consensus that microbial communities in fecal samples do not represent the whole GI microbiota. Studies in humans suggest that microbial communities in the duodenum and colon are not represented by those in feces because of the large differences in microbial profiles [31], and these studies emphasized the need to examine tissue biopsies in addition to fecal samples [5], proposing that standard forceps mucosal biopsy samples can represent bacterial populations [4]. Compared with human studies, studies in other mammals are more comprehensive because a larger number of gut segments can be involved in the analyses. Several studies in mice [25, 32] support the utility of fecal samples for studying the gut microbiota, because microbial communities in fecal samples were observed to be similar to those in the lower GI tract, which is supported by studies conducted in rhesus macaques [30], pigs [33] and equines [34].

Compared with previous studies, the strength of the current study lies in the following: 1) it involved the use of gut segments from the upper GI tract to the lower GI tract and feces, providing a relatively comprehensive view of the spatial relationships of the gut microbiota; 2) the microbial relationships were partitioned into two parts, i.e., microbial community membership and structure, providing multiangle observations to identify microbial relationships between feces and the GI tract; and 3) a massive number of individuals was sampled, which is significant for investigations of gut spatial relationships, as the sizes of most of the above studies did not exceed twenty. The considerable sample size would provide more comprehensive insights into exploring the utility of fecal samples in studies of the gut microbiota.

Because of the specific and significant roles in nutrition and health [17, 35], ceca have been widely investigated in birds [36, 37], especially chickens [20, 21, 38]. *Bacteroides* was observed as the dominant taxa in our study (Additional file 3: Figure S1D) and in most other studies [39, 40], although some reports observed a predominance of *Clostridiales* members in ceca [14, 41]. Although the cecal microbial community may sometimes be linked to diet [36], the nearly consistent results across studies suggests that the cecal microbial community is stable. This finding might be due to ceca having a special blind-ended structure and being located in the lower GI tract, providing a stable and anaerobic environment for microbes and longer storage periods of the contents, in contrast to the rapid transit environment in the small intestine [42]. In addition to the microbial composition, Stanley et al. [22] also compared microbial differences and similarities between ceca and feces in chicken. They observed that 88.55% of all OTUs, containing 99.25% of all sequences, were shared by the ceca and feces, similar to the observations in the current study. These results indicate that except for some rare microbial members, most microbes in the ceca can be detected in fecal samples.

The microbial relationships between the ceca and small intestine have been rarely reported in birds. Choi et al. [14] compared the percentage of shared OTUs among ceca and three small intestinal sections but observed low percentages between segments (ranging from 1.2 to 2.9%, representing from 38.7 to 65.5% of sequences). The percentages reported in another study (60.2% for the duodenum, 50.5% for the jejunum and 43.5% for the ileum, which were calculated from Figure 3 in their article) were higher than those in Choi’s study. In contrast, the results of Xiao’s study presented an opposite trend from our findings, i.e., the percentages of shared OTUs in Xiao’s study decreased from the duodenum to the jejunum and ileum, demonstrating a reversed-anatomical feature compared with the current study. These inconsistent results might be attributable to differences among species, diets or other environmental factors, but the small sample size in Xiao’s study may be an important reason for these inconsistencies.

## Conclusion

Overall, we assessed the efficacy of using fecal samples to represent GI microbiota in birds and analyzed potential factors affecting this efficacy. With highly shared microbial members, fecal samples have the good potential to be used to detect most microbial species in the small intestine and ceca with gut anatomical features. However, analyses of microbial structures using fecal samples as the proxy for the gut in longitudinal microbial studies should be interpreted with caution. This study attempts to identify the microbial relationships between feces and the intestine in birds, which will help extend our understanding of the bird gut microbiota and provide future directions regarding the usage of fecal samples in studies of the gut microbiome.

## Methods

### Animal model

The complete procedure was performed according to the guidelines established by the Animal Care and Use Committee of China Agricultural University (permit number: SYXK 2013-0013).

The slow-growing yellow broiler was used as the animal model in this study, and the birds were obtained from Wen’s Nanfang Poultry Breeding Co., Ltd. in Guangdong Province of China. Two hundred and six birds with similar body weights were selected and raised on the ground with *ad libitum* feeding and nipple drinkers. The birds were fed a common maize-soybean-based diet throughout the duration of the experiment. No antibiotics were applied during the thirty-five days before sample collection. Because chickens are the largest population of birds on earth, the chicken was selected as a bird model for this investigation. The slow-growing yellow broiler has not been highly selected for production, making this breed of chicken closer to the ancestral birds.

### Sample collection

Fresh fecal samples were collected from each bird as soon as excreta was discharged through the cloaca at 77 days of age with the average body weight was 2.32 kg. Next, all the birds were humanely euthanized by cervical dislocation and subsequently dissected. The contents and mucosal surfaces of the duodenum, jejunum, ileum and cecum were collected immediately after dissection. To ensure the consistency of samples among individuals, a 10-cm-long fixed section of the duodenum and jejunum, the whole ileum and a pair of ceca were selected for sampling from each bird. The contents and mucosa were mixed uniformly before collection. All samples were immediately placed in liquid nitrogen and then stored at −80°C. Both the intestinal contents and mucosa were sampled based on the consideration that the microbes from both sources may contribute to host interactions with respect to nutrient metabolism and immunity [43].

### DNA extraction and 16S rRNA gene sequencing

DNA was extracted from intestinal and fecal samples using a QIAamp DNA stool mini kit (QIAGEN, cat#51504) [44] following the manufacturer’s instructions. PCR amplification of the V4 region of the bacterial 16S rRNA gene was performed using the forward primer 520F (5’-AYTGGGYDTAAAGNG-3’) and the reverse primer 802R (5’-TACNVGGGTATCTAATCC-3’). Sample-specific 7-bp barcodes were incorporated into the primers for multiplex sequencing. The PCR reactions contained 5 μl of Q5 reaction buffer (5×), 5 μl of Q5 High-Fidelity GC buffer (5×), 0.25 μl of Q5 High-Fidelity DNA Polymerase (5 U/μl), 2 μl of dNTPs (2.5 mM), 1 μl (10 µM) of each forward and reverse primer, 2 μl of DNA template, and 8.75 μl of ddH_2_O. Thermal cycling consisted of initial denaturation at 98°C for 2 min, followed by 25 cycles of denaturation at 98°C for 15 s, annealing at 55°C for 30 s, and extension at 72°C for 30 s, with a final extension at 72°C for 5 min. PCR amplicons were purified using Agencourt AMPure Beads (Beckman Coulter, Indianapolis, IN) and quantified using a PicoGreen dsDNA Assay kit (Invitrogen, Carlsbad, CA, USA). After the quantification step, amplicons were pooled in equal amounts, and 2 × 300 bp paired-end sequencing was performed using an Illumina MiSeq platform with the MiSeq Reagent kit v3 at Shanghai Personal Biotechnology Co., Ltd. (Shanghai, China). The raw data on which the conclusions of the manuscript rely have been deposited in the National Center for Biotechnology Information (NCBI) database (accession numbers SRP139192, SRP139193 and SRP139195).

### Analysis of sequencing data

Data analysis was performed using the Quantitative Insights Into Microbial Ecology (QIIME, v1.8.0) pipeline [45], due to the advantages of QIIME [46-48]. Briefly, raw sequencing reads with exact matches to the barcodes were assigned to respective samples and identified as valid sequences. The low-quality sequences were filtered based on the following criteria [49, 50]: length < 150 bp, average Phred score < 20, ambiguous bases, and mononucleotide repeats > 8 bp. Paired-end reads were assembled using FLASH [51], and chimera detection was performed with QIIME. After quality control, four fecal samples were excluded due to low sequence quality that was potentially caused by a technical artifact. The remaining high-quality sequences were clustered into operational taxonomic units (OTUs) at a 97% sequence identity using an open-reference OTU picking protocol against the Silva database (SILVA128) [52-54].

We focused on open-reference OTU picking for these analyses because this method yields substantially more taxonomic identifications with sequences that failed to hit the reference database than do closed-reference methods [55]. The open-reference method can provide more information for comparisons among intestinal segments or feces. The singleton OTUs were discarded because such OTUs can occur due to sequencing errors. Only OTUs representing more than 0.001% of the total filtered OTUs were retained to improve the efficiency of the analysis. Because the sequencing and sampling quantity varied among individuals, we rarefied the data to the lowest numbers of sequences per sample to control for sampling effort in diversity analyses. Alpha and beta diversity of individual OTUs were calculated with postrarefaction data and the phylogenetic tree. Principal coordinate analysis (PCoA) was performed using the unweighted or weighted UniFrac distance [56] for different intestinal segments and feces. To decrease the data noise, only OTUs that were present in more than 3 samples at each sampling site were used to analyze the effect of microbial membership. The correlations between the mean fecal and segmental genera abundance were calculated using the method described in a study of rhesus macaques [30]. These methods could primarily provide the number and diversity of microbes in feces and each segment which would help to quantitatively understand the relationships of microbial communities between feces and GI tract.

### Statistical analysis

Venn plots were generated for intestinal segment or feces samples at the OTU level using the VennDiagram package in **R**. Spearman correlation analysis was performed in package psych in **R**. Paired Student’s *t*-test was used to compare the microbial UniFrac distance between two sampling sites. Mann-Whitney test was performed to identify the differences of each genus between two sampling sites.

## Abbreviations

GI: Gastrointestinal
FD, FJ, FI and FC: The unweighted or weighted UniFrac distance between feces and duodenum, jejunum, ileum or cecum, respectively
NGS: Next-generation sequencing
OTU: Operational taxonomic unit.

## Declarations

### Consent for publication

Not applicable

### Availability of data and material

The raw data on which the conclusions of the manuscript rely has been deposited in the National Center for Biotechnology Information (NCBI) database (accession number SRP139192, SRP139193 and SRP139195).

### Authors’ contributions

WY, JXZ and NY designed the study. WY, JXZ, CLW, CLJ, DXZ, YHC and CJS collected the samples. WY analyzed the data and wrote the manuscript. CLW assisted in construction of the figures. CJS and NY assisted in data analyzing and contributed to the revisions. All authors read and approved the final manuscript.

## Acknowledgements

We thank Prof. Guiyun Xu for demonstration of dissection; Guangqi Li, Zhongyi Duan, Shanshan Xie, Jingwei Yuan, Dehe Wang, Zebin Zhang, Xingzheng Li, Yajie Li, Chunning Mai and Zhenfei Jiang for assistance with the sample collection; and Dr. Zhengsheng Xue for advice on the study. The current study was funded in part by Programs for Changjiang Scholars and Innovative Research in University (IRT_15R62) and China Agriculture Research System (CARS-40).

## References

1. Rosenbaum M, Knight R, Leibel RL: The gut microbiota in human energy homeostasis and obesity. Trends Endocrinol Metab. 2015; 26:493–501.

2. Shin NR, Lee JC, Lee HY, Kim MS, Whon TW, Lee MS, Bae JW: An increase in the Akkermansia spp. population induced by metformin treatment improves glucose homeostasis in diet-induced obese mice. Gut. 2014; 63:727–35.

3. Brisbin JT, Gong J, Sharif S: Interactions between commensal bacteria and the gut-associated immune system of the chicken. Anim Health Res Rev. 2008; 9:101–10.

4. Lavelle A, Lennon G, O’sullivan O, Docherty N, Balfe A, Maguire A, Mulcahy HE, Doherty G, O’Donoghue D, Hyland J et al: Spatial variation of the colonic microbiota in patients with ulcerative colitis and control volunteers. Gut. 2015; 64:1553–61.

5. Gevers D, Kugathasan S, Denson LA, Vázquezbaeza Y, Van TW, Ren B, Schwager E, Knights D, Song SJ, Yassour M: The treatment-naive microbiome in new-onset Crohn’s disease. Cell Host Microbe. 2014; 15:382–92.

6. McCormack UM, Curiã O T, Buzoianu SG, Prieto ML, Ryan T, Varley P, Crispie F, Magowan E, Metzler-Zebeli BU, Berry D: Exploring a possible link between the intestinal microbiota and feed efficiency in pigs. Appl Environ Microbiol. 2017; 83: e00380–17.

7. Johnson TJ, Youmans BP, Noll S, Cardona C, Evans NP, Karnezos TP, Ngunjiri JM, Abundo MC, Lee CW: A consistent and predictable commercial broiler chicken bacterial microbiota in antibiotic-free production displays strong correlations with performance. Appl Environ Microbiol. 2018; 84:e00362–18.

8. Dumonceaux TJ, Hill JE, Hemmingsen SM, Van Kessel AG: Characterization of intestinal microbiota and response to dietary virginiamycin supplementation in the broiler chicken. Appl Environ Microbiol. 2006; 72:2815–23.

9. Yeoman CJ, Chia N, Jeraldo P, Sipos M, Goldenfeld ND, White BA: The microbiome of the chicken gastrointestinal tract. Anim Health Res Rev. 2012; 13:89–99.

10. Stanley D, Hughes RJ, Moore RJ: Microbiota of the chicken gastrointestinal tract: influence on health, productivity and disease. Appl Microbiol Biotechnol. 2014; 98:4301–10.

11. Shaufi MAM, Sieo CC, Chong CW, Gan HM, Ho YW: Deciphering chicken gut microbial dynamics based on high-throughput 16S rRNA metagenomics analyses. Gut Pathog. 2015; 7:4.

12. Clavijo V, Mjv F: The gastrointestinal microbiome and its association with the control of pathogens in broiler chicken production: A review. Poult Sci. 2018; 97:1006–21.

13. Gong J, Si W, Forster R, Huang R, Yu H, Yin Y, Yang CY: 16S rRNA gene-based analysis of mucosa-associated bacterial community and phylogeny in the chicken gastrointestinal tracts: from crops to ceca. FEMS Microbiol Ecol. 2007; 59:147–57.

14. Choi JH, Kim GB, Cha CJ: Spatial heterogeneity and stability of bacterial community in the gastrointestinal tracts of broiler chickens. Poult Sci. 2014; 93:1942–50.

15. Xiao Y, Xiang Y, Zhou W, Chen J, Li K, Yang H: Microbial community mapping in intestinal tract of broiler chicken. Poult Sci. 2016; 96:1387–93.

16. Mead GC: Microbes of the avian cecum: types present and substrates utilized. J Exp Zool Suppl. 1989; 252:48–54.

17. Clench MH, Mathias JR: The avian cecum: A review. Wilson Bulletin. 1995; 107:93–121.

18. Obst BS, Diamond JM: Interspecific variation in sugar and amino acid transport by the avian cecum. J Exp Zool Suppl. 1989; 252:117–26.

19. Gasaway WC, White RG, Dan FH: Digestion of dry matter and absorption of water in the intestine and cecum of rock ptarmigan. Condor. 1976; 78:77–84.

20. Stanley D, Geier MS, Hughes RJ, Denman SE, Moore RJ: Highly variable microbiota development in the chicken gastrointestinal tract. PLOS One. 2013; 8:e84290.

21. Sergeant MJ, Constantinidou C, Cogan TA, Bedford MR, Penn CW, Pallen MJ: Extensive microbial and functional diversity within the chicken cecal microbiome. PLOS One. 2014; 9:e91941.

22. Stanley D, Geier MS, Chen H, Hughes RJ, Moore RJ: Comparison of fecal and cecal microbiotas reveals qualitative similarities but quantitative differences. BMC Microbiol. 2015; 15:51.

23. Rizzo JM, Buck MJ: Key principles and clinical applications of “next-generation” DNA sequencing. Cancer Prev Res. 2012; 5:887–900.

24. Daber R, Sukhadia S, Morrissette JJ: Understanding the limitations of next generation sequencing informatics, an approach to clinical pipeline validation using artificial data sets. Cancer Genet. 2013; 206:441–8.

25. Suzuki TA, Nachman MW: Spatial heterogeneity of gut microbial composition along the gastrointestinal tract in natural populations of house mice. PLOS One. 2016; 11:e163720.

26. Sekelja M, Rud I, Knutsen SH, Denstadli V, Westereng B, Næs T, Rudi K: Abrupt temporal fluctuations in the chicken fecal microbiota are explained by its gastrointestinal origin. Appl Environ Microbiol. 2012; 78:2941–8.

27. Gu S, Chen D, Zhang JN, Lv X, Wang K, Duan LP, Nie Y, Wu XL: Bacterial community mapping of the mouse gastrointestinal tract. PLOS One. 2013; 8:e74957.

28. Dias J, Marcondes MI, Motta DSS, Cardoso DMB, Noronha MF, Resende RT, Machado FS, Mantovani HC, Dill-Mcfarland KA, Suen G: Bacterial community dynamics across the gastrointestinal tracts of dairy calves during preweaning development. Appl Environ Microbiol. 2018; 84:e02675–17.

29. Yan W, Sun C, Yuan J, Yang N: Gut metagenomic analysis reveals prominent roles of Lactobacillus and cecal microbiota in chicken feed efficiency. Sci Rep. 2017; 7:45308.

30. Yasuda, Koji, Keunyoung, Ren, Boyu, Nbsp T, Franzosa: Biogeography of the intestinal mucosal and lumenal microbiome in the rhesus macaque. Cell Host Microbe. 2015; 17:385–91.

31. Stearns JC, Lynch MDJ, Senadheera DB, Tenenbaum HC, Goldberg MB, Cvitkovitch DG, Croitoru K, Morenohagelsieb G, Neufeld JD: Bacterial biogeography of the human digestive tract. Sci Rep. 2011; 1:170.

32. Li D, Chen H, Mao B, Yang Q, Zhao J, Gu Z, Zhang H, Chen YQ, Chen W: Microbial biogeography and core microbiota of the rat digestive tract. Sci Rep. 2017; 8:45840.

33. Zhao W, Wang Y, Liu S, Huang J, Zhai Z, He C, Ding J, Wang J, Wang H, Fan W et al: The dynamic distribution of porcine microbiota across different ages and gastrointestinal tract segments. PLoS One. 2015; 10:e117441.

34. Ericsson AC, Johnson PJ, Lopes MA, Perry SC, Lanter HR: A microbiological map of the healthy equine gastrointestinal tract. PLoS One. 2016; 11:e166523.

35. Waite DW, Taylor MW: Characterizing the avian gut microbiota: membership, driving influences, and potential function. Front Microbiol. 2014; 5:223.

36. Wienemann T, Schmitt-Wagner D, Meuser K, Segelbacher G, Schink B, Brune A, Berthold P: The bacterial microbiota in the ceca of Capercaillie (Tetrao urogallus) differs between wild and captive birds. Syst Appl Microbiol. 2011; 34:542–51.

37. Matsui H, Kato Y, Chikaraishi T, Moritani M, Bantokuda T, Wakita M: Microbial diversity in ostrich ceca as revealed by 16S ribosomal RNA gene clone library and detection of novel Fibrobacter species. Anaerobe. 2010; 16:83–93.

38. Oakley BB, Lillehoj HS, Kogut MH, Kim WK, Maurer JJ, Pedroso A, Lee MD, Collett SR, Johnson TJ, Cox NA: The chicken gastrointestinal microbiome. FEMS Microbiol. Lett. 2014; 360:100–12.

39. Wei S, Morrison M, Yu Z: Bacterial census of poultry intestinal microbiome. Poult Sci. 2013; 92:671–83.

40. Tillman GE, Haas GJ, Wise MG, Oakley B, Smith MA, Siragusa GR: Chicken intestine microbiota following the administration of lupulone, a hop-based antimicrobial. FEMS Microbiol Ecol. 2011; 77:395–403.

41. Cressman MD, Yu Z, Nelson MC, Moeller SJ, Lilburn MS, Zerby HN: Interrelations between the microbiotas in the litter and in the intestines of commercial broiler chickens. Appl Environ Microbiol. 2010; 76:6572–82.

42. Clench MH, Mathias JR: A complex avian intestinal motility response to fasting. Am J Physiol. 1992; 262:498–504.

43. Smith CC, Snowberg LK, Gregory CJ, Knight R, Bolnick DI: Dietary input of microbes and host genetic variation shape among-population differences in stickleback gut microbiota. ISME J. 2015; 9:2515–26.

44. Zhao LL, Wang G, Siegel P, He C, Wang HZ, Zhao WJ, Zhai ZX, Tian FW, Zhao JX, Zhang H et al: Quantitative genetic background of the host influences gut microbiomes in chickens. Sci Rep. 2013; 3:1163.

45. Caporaso JG, Kuczynski J, Stombaugh J, Bittinger K, Bushman FD, Costello EK, Fierer N, Peña AG, Goodrich JK, Gordon JI: QIIME allows analysis of high-throughput community sequencing data. Nat Methods. 2010; 7:335–6.

46. Nilakanta H, Drews KL, Firrell S, Foulkes MA, Jablonski KA: A review of software for analyzing molecular sequences. BMC Res Notes. 2014; 7:830.

47. Plummer E, Twin J: A comparison of three bioinformatics pipelines for the analysis of preterm gut microbiota using 16S rRNA gene sequencing data. J Proteomics Bioinformatics. 2015; 8:283–91.

48. Pollock J, Glendinning L, Wisedchanwet T, Watson M: The madness of microbiome: attempting to find consensus “best practice” for 16S microbiome studies. Appl Environ Microbiol. 2018; 84:e02627–17.

49. Gill SR, Pop M, Deboy RT, Eckburg PB, Turnbaugh PJ, Samuel BS, Gordon JI, Relman DA, Fraserliggett CM, Nelson KE: Metagenomic analysis of the human distal gut microbiome. Science. 2006; 312:1355–9.

50. Chen H, Jiang W: Application of high-throughput sequencing in understanding human oral microbiome related with health and disease. Front Microbiol. 2014; 5:508.

51. Magoč T, Salzberg SL: FLASH: fast length adjustment of short reads to improve genome assemblies. Bioinformatics. 2011; 27:2957–63.

52. Quast C, Pruesse E, Yilmaz P, Gerken J, Schweer T, Yarza P, Peplies J, Glöckner FO: The SILVA ribosomal RNA gene database project: improved data processing and web-based tools. Nucleic Acids Res. 2013; 41:D590–6.

53. Yilmaz P, Parfrey L, Yarza P, Gerken J, Pruesse E, Quast C, Schweer T, Peplies J, Ludwig W, Glöckner F: The SILVA and “All-species Living Tree Project (LTP)” taxonomic frameworks. Nucleic Acids Res. 2014; 42:D643–8.

54. Balvočiūtė M, Huson DH: SILVA, RDP, Greengenes, NCBI and OTT — how do these taxonomies compare? BMC Genomics. 2017; 18(S2):114.

55. Rideout JR, He Y, Navasmolina JA, Walters WA, Ursell LK, Gibbons SM, Chase J, Mcdonald D, Gonzalez A, Robbinspianka A: Subsampled open-reference clustering creates consistent, comprehensive OTU definitions and scales to billions of sequences. PeerJ. 2014; 2:e545.

56. Lozupone C, Knight R: UniFrac: a new phylogenetic method for comparing microbial communities. Appl Environ Microbiol. 2005; 71:8228–35.

